# Mobile-CRISPRi: Enabling Genetic Analysis of Diverse Bacteria

**DOI:** 10.1101/315499

**Authors:** Jason M. Peters, Byoung-Mo Koo, Ramiro Patino, Gary E. Heussler, Cameron C. Hearne, Yuki F. Inclan, John S. Hawkins, Candy H. S. Lu, M. Michael Harden, Hendrik Osadnik, Joseph E. Peters, Joanne N. Engel, Rachel J. Dutton, Alan D. Grossman, Carol A. Gross, Oren S. Rosenberg

## Abstract

The vast majority of bacteria, including human pathogens and microbiome species, lack genetic tools needed to systematically associate genes with phenotypes. This is the major impediment to understanding the fundamental contributions of genes and gene networks to bacterial physiology and human health. CRISPRi, a versatile method of blocking gene expression using a catalytically inactive Cas9 protein (dCas9) and programmable single guide RNAs (sgRNAs), has emerged as a powerful genetic tool to dissect the functions of essential and non-essential genes in species ranging from bacteria to human. However, the difficulty of establishing effective CRISPRi systems in non-model bacteria is a major barrier to its widespread use to dissect bacterial gene function. Here, we establish “Mobile-CRISPRi”, a suite of CRISPRi systems that combine modularity, stable genomic integration and ease of transfer to diverse bacteria by conjugation. Focusing predominantly on human pathogens associated with antibiotic resistance, we demonstrate the efficacy of Mobile-CRISPRi in Proteobacteria and Firmicutes at the individual gene scale by examining drug-gene synergies and at the library scale by systematically phenotyping conditionally essential genes involved in amino acid biosynthesis. Mobile-CRISPRi enables genetic dissection of non-model bacteria, facilitating analyses of microbiome function, antibiotic resistances and sensitivities, and comprehensive screens for host-microbe interactions.

## Main text

CRISPRi (Clustered Regularly Interspaced Short Palindromic Repeats interference) is a programmable method for controlling gene expression that has enabled systematic interrogation of gene phenotypes in diverse organisms^1–6^. In bacterial CRISPRi, an sgRNA-dCas9 complex binds to a target gene by base-pairing and reduces gene expression by sterically blocking transcription elongation (Fig. 1a)^1,4^. New CRISPRi targets are easily programmed by substituting the first 20 nt of the sgRNA sequence (spacer) to match the non-template strand of the target gene, making design and construction of CRISPRi libraries that target specific sets of genes or the entire genome straightforward^4,7,8^. Genetic screens using CRISPRi libraries have contributed new insights into fundamental biology and molecular medicine including identifying functions for uncharacterized essential genes^4,7^ and drug modes of action^4,9^.

**Fig. 1.**
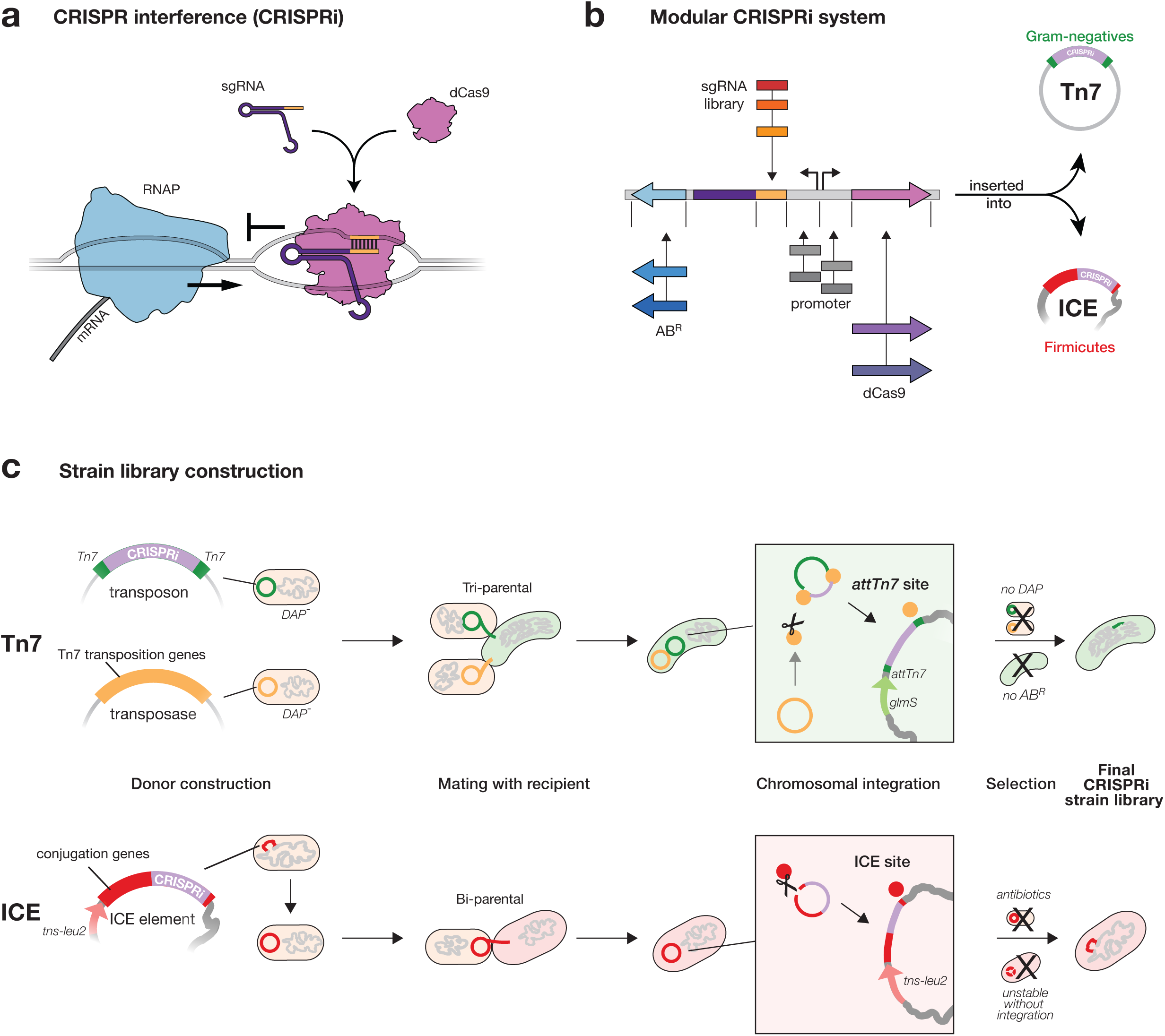
Mobile-CRISPRi overview. **a**, Mechanism of CRISPRi repression. A dCas9-sgRNA complex binds to DNA by base-pairing and sterically blocks progression of RNA polymerase (RNAP), reducing gene expression. **b,** Mobile-CRISPRi modularity. Individual modules are flanked by unique restriction sites, which can be used for cloning sgRNA libraries or exchanging other components (e.g. antibiotic resistance marker (AB^R^), or *dcas9* promoter). **c,** Strain construction using Mobile-CRISPRi. Top: a Tn*7* transposon carrying CRISPRi components (shown in b) and a plasmid containing Tn*7* transposition genes are transferred to recipient bacteria by tri-parental mating. Donor cells contain a chromosomal copy of the RP4 transfer machinery used to mobilize the Tn*7* plasmids. Once inside the recipient cell, Tn*7* transposition proteins integrate the CRISPRi DNA (purple) flanked by left and right Tn*7* end sequences (green) into the recipient genome downstream of the *glmS* gene. Selection on antibiotic plates lacking DAP eliminates the *E. coli* donors and retains recipients with an integrated CRISPRi system. Bottom: An ICE element carrying CRISPRi components is transferred to recipient bacteria by bi-parental mating. Once inside the recipient cell, the ICE integrase inserts ICE into *trnS-leu2*. Double antibiotic plates that select for ICE and for the intrinsic resistance of the recipient strain (Streptomycin-resistance in this work) are used to identify recipients with an integrated CRISPRi system.

CRISPRi provides several advantages over other methods for genetic manipulation in bacteria. CRISPRi knockdowns can be induced^1, 3–6^ and titrated/tuned^4,10^, enabling depletion of essential gene products without complex strain construction strategies that remove genes from their native regulation. Dissecting genetic redundancy via multiplexed CRISPRi targeting several genes in the same cell^4,11^ requires markedly less effort than construction of multiple-deletion strains. At the genome scale, CRISPRi expands on prior transposon-based gene perturbation methods such as Tn-seq^12^ by allowing all genes—including essential genes that cannot be studied through deletion—to be systematically targeted so that a relatively small strain library provides comprehensive coverage of the genome. Moreover, the DNA sequences encoding sgRNAs serve as unique barcodes to differentiate CRISPRi strains mixed in a pool, allowing for competitive fitness measurements using next generation sequencing^8^. CRISPRi blocks expression of downstream genes in operons^1,4^, but this property can be used to further simplify libraries by targeting operons instead of genes.

Despite these advantages, CRISPRi has been used in only a few bacterial species both because CRISPRi has been transferred using species-specific^4^ or narrow host-range^1,3,5,6,13^ strategies, and because components need to be optimized for function in different species. To overcome this barrier, we developed “Mobile-CRISPRi”—a suite of modular and transferable CRISPRi components that can stably integrate into the genomes of diverse bacteria. The modularity of every component of Mobile-CRISPRi makes it straightforward to clone in organism-specific sgRNA libraries and other components (e.g. promoters, Fig. 1b). Mobile-CRISPRi achieves transfer and genomic integration by distinct mechanisms for Proteobacteria and Firmicutes. For Proteobacteria, Mobile-CRISPRi is transferred from *Escherichia coli* using the broad host range RP4 plasmid conjugation machinery, and is orientation and site specifically integrated into the recipient genome downstream of the highly conserved *glmS* gene using the extensively characterized Tn*7* transposition system (Fig. 1c, top)^14,15^. Consistent with previous reports^15^, this strategy was unsuccessful in Firmicutes such as *Bacillus subtilis* (Supplementary Fig. 1), leading us to develop a strategy to transfer CRISPRi using the ICE *Bs1* conjugation and integration machinery. Here, Mobile-CRISPRi is transferred from *B. subtilis* to other Bacillales Firmicutes (*e.g.*, *Staphylococcus aureus*—although the host range has not been experimentally defined), and integrated into *trnS-leu2*^16^(Fig. 1c, bottom). Critically, Mobile-CRISPRi integrations downstream of *glmS* and into *trnS-leu2* do not disrupt those gene’s functions^14,16^, occur in a specified orientation, and are stable in the absence of selection for ≥ 50 generations (Fig. 2a), enabling studies of antibiotic function in which maintaining selection is problematic or impossible.

**Fig. 2.**
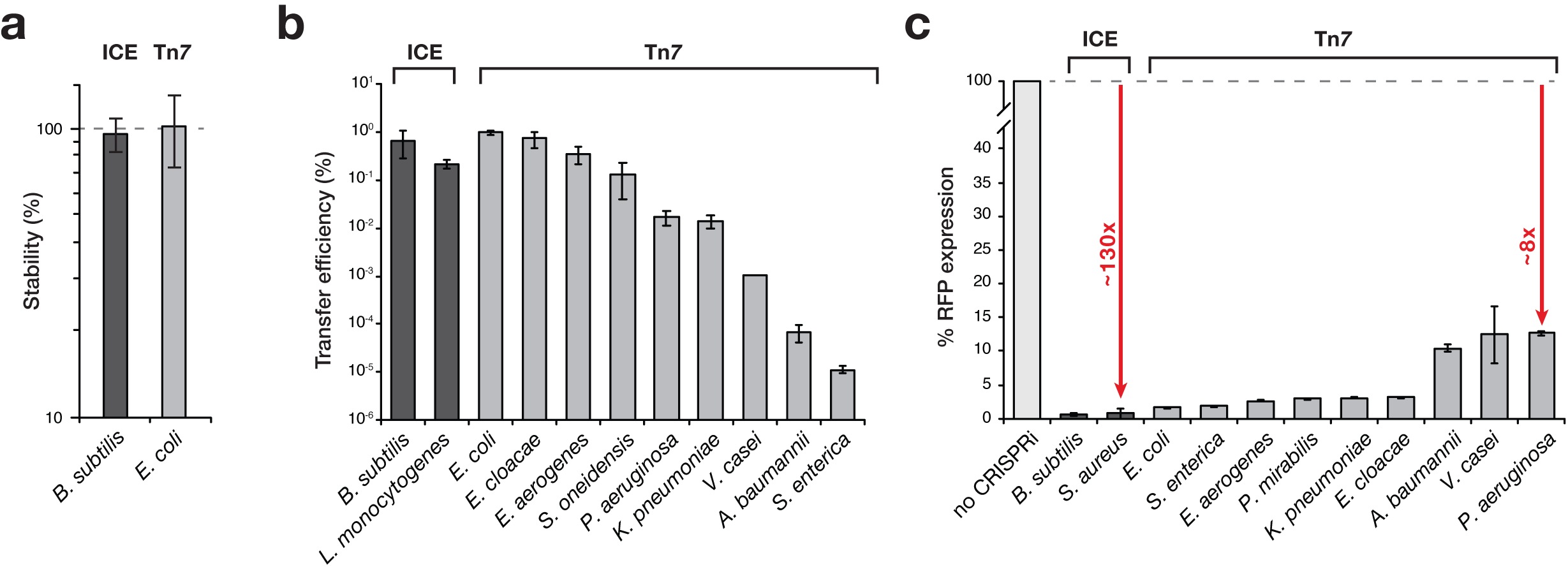
Mobile-CRISPRi stability, transfer, and knockdown efficiency. **a,** Mobile-CRISPRi stability after 50 generations of growth in the absence of antibiotic selection. Stability is the plating efficiency on kanamycin (the marker associated with Mobile-CRISPRi) vs no antibiotic. **b,** Mobile-CRISPRi transfer and integration efficiency. ICE or Tn*7* containing CRISPRi was transferred to the species listed. Efficiency was calculated as: %AB^R^/total recipients. **c,** Efficiency of Mobile-CRISPRi knockdown in various species. Knockdown was tested using a Mobile-CRISPRi variant containing a constitutively expressed *rfp* reporter and an sgRNA targeting *rfp*. RFP expression was normalized to a strain lacking either *dcas9* (for *P. aeruginosa*) or an sgRNA (all others). Data are represented as mean ± s.d. The full species names for the strains used in panels b and c and their corresponding *rfp* fold knockdowns are: *Bacillus subtilis* (182-fold), *Listeria monocytogenes* (ND), *Escherichia coli* (65-fold), *Enterobacter cloacae* (32-fold), *Enterobacter aerogenes* (40-fold), *Shewanella oneidensis* (ND), *Pseudomonas aeruginosa* (8-fold), *Klebsiella pneumoniae* (34-fold), *Vibrio casei* (8-fold), *Acinetobacter baumannii* (10-fold), *Salmonella enterica* (54-fold), *Staphylococcus aureus* (132-fold), and *Proteus mirabilis* (35-fold).

To assess the efficacy of Mobile-CRISPRi in non-model bacteria, we measured CRISPRi transfer and knockdown efficiency primarily in species involved in human disease. We quantified transfer efficiency by counting the number of recipient colonies (i.e., transconjugants) on selective agar plates (Fig. 2b). We found that most strains showed transfer efficiencies that were sufficient for genome-scale sgRNA library construction (e.g., *Enterobacter sp.* ~10^-2^-10^-3^, and *L. monocytogenes* ~10^-2^), whereas other strains were more suited for single gene knockdown approaches (e.g., *Acinetobacter baumanii* ~10^-6^).

To assess CRISPRi knockdown efficacy across species, we created a “test” version of Mobile-CRISPRi consisting of *rfp* (encoding Red Fluorescent Protein, or RFP) and either an sgRNA targeting *rfp*, or a “control” version lacking an sgRNA to normalize *rfp* expression. Our quantification of *rfp* knockdown in single cells using flow cytometry indicated that knockdown efficiency ranged from ~8-fold in *Pseudomonas aeruginosa* to ~130-fold in *S. aureus*, with a median knockdown of ~35-fold across all measured species (Fig. 2c). We confirmed that CRISPRi was also functional against native genes by targeting *P. aeruginosa* pyocyanin production (Supplementary Fig. 2). In this visual assay, the culture appears yellow, as production of the blue pigment pycocyanin is decreased.

Our initial assessment of Mobile-CRISPRi used pathogenic strains possessing at least rudimentary tools for perturbing gene function. To determine whether Mobile-CRISPRi functions in an environmental isolate with no existing genetic system, we tested transfer and knockdown in *Vibrio casei*, a γ-proteobacterium originally isolated from French wash-rind cheeses^17^ and broadly associated with cheese microbiomes^18^. We found that Mobile-CRISPRi transferred to *V. casei* with library scale efficiency (~10^-3^, Fig. 2a), and a modest, but useful knockdown efficiency (~8-fold, Fig. 2b). The modular nature of Mobile-CRISPRi allows for further optimization of knockdown efficiency; for instance, by using *Vibrio*-specific promoters for *dcas9* and sgRNA expression. We conclude that Mobile-CRISPRi is an effective genetic tool for gene knockdowns in diverse, non-model bacteria.

The emergence of multi-drug resistant pathogenic bacteria is an urgent threat to human health that requires both new antibiotics and a better understanding how existing antibiotics work^19^. Knowledge of the mechanism by which antibiotics kill bacteria—the mode of action (MOA)—is critical to advance new antibiotics from the laboratory to the clinic^20^. Because the full complement of genes in a bacterial genome (i.e., genetic background) can affect antibiotic function^20^, the MOA should ideally be determined directly in clinically relevant strains. However, most pathogenic bacterial lack genetic tools to systematically perturb the functions of essential genes that typically encode antibiotic targets. Importantly, we previously showed that the ability to titrate the knockdown level enables the systematic study of essential genes in *B subtilis*. A low (~3-fold) level of knockdown allowed sufficient growth to determine the MOA of an uncharacterized antibiotic by virtue of its synergistic effects on growth^4^.

To explore whether Mobile-CRISPRi could be used to explore MOA in pathogenic Proteobacteria associated with antibiotic resistance (i.e., Gram-negative rods), we examined synergy between the antibiotic trimethoprim and the essential gene *folA*, which encodes the trimethoprim target dihydrofolate reductase^21^. We found that targeting *folA* with CRISPRi in *Enterobacter aerogenes*, *Klebsiella pneumoniae*, and *P. aeruginosa* strongly increased sensitivity to trimethoprim (Fig. 3) and shifted the minimal inhibitory concentration (MIC) by 2-4-fold, depending on the species (Fig. 3a). Even though CRISPRi knockdown in *P. aeruginosa* is at a lower efficiency compared to other strains (Fig. 2b), there was still a clear shift toward sensitivity (Fig. 3a). Moreover, concentrations of trimethoprim below the MIC for the WT, completely inhibited growth of the *folA* knockdown strains, clearly demonstrating synergy (Fig. 3b). Thus, Mobile-CRISPRi targeting essential genes can be used to generate sensitized strains for antibiotic MOA studies.

**Fig. 3.**
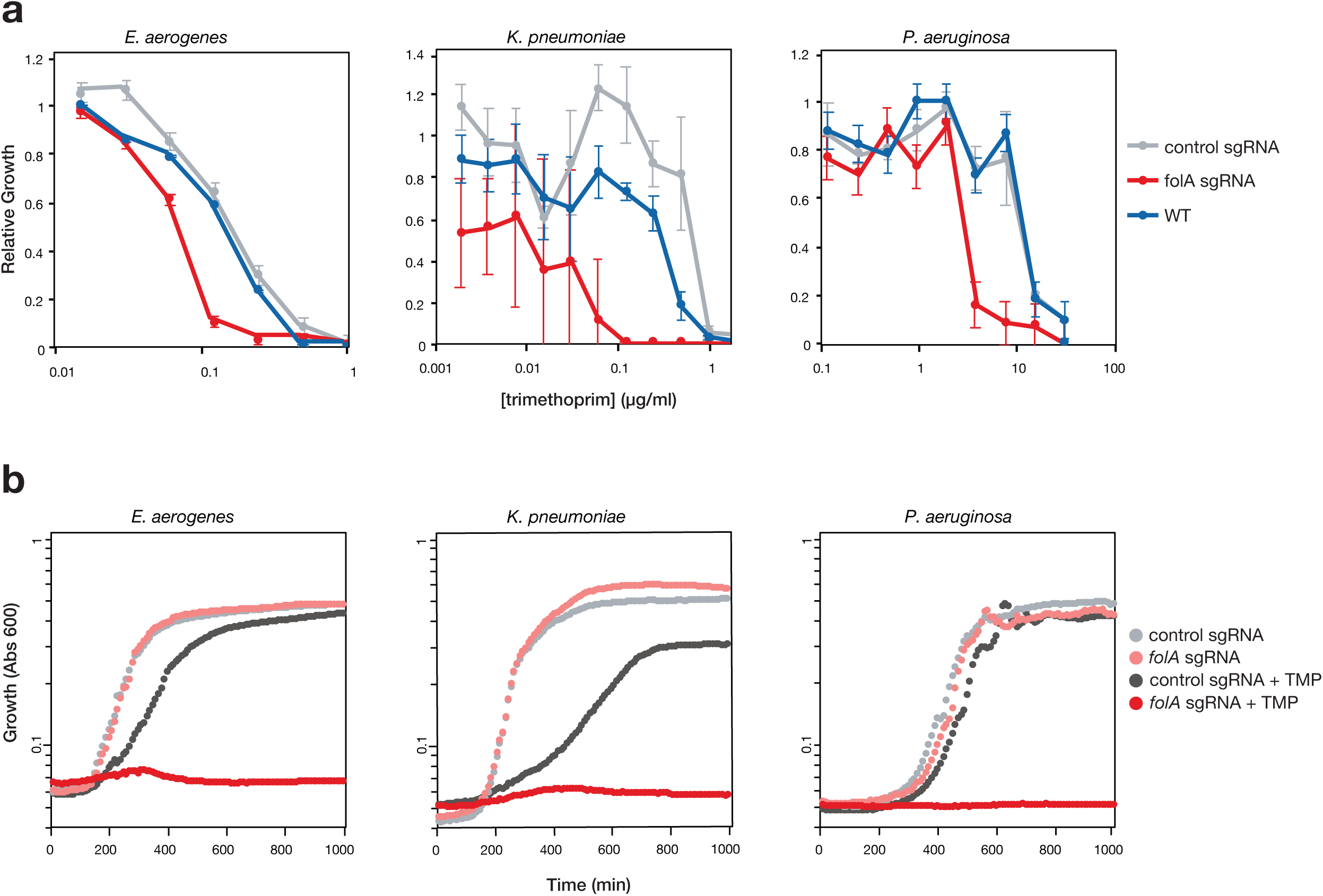
CRISPRi knockdown of *folA* increases sensitivity to trimethoprim in multiple species. **a**, MIC assays for trimethoprim sensitivity in *P. aeruginosa* and *E. aerogenes* with or without Mobile-CRISPRi targeting *folA*. *K. pneumoniae* clumping increased the measurement error, although sensitivity is readily apparent. Data are represented as mean ± s.d. **b**, Mobile-CRISPRi targeting *folA* or with a non-targeting control guide in various species were grown with or without trimethoprim (0.125, 0.5, and 8 μg/ml trimethoprim for *E. areogenes*, *K. pneumoniae*, and *P. aeruginosa*, respectively), and with partial induction of CRISPRi (100 μM IPTG for *E. areogenes*, *K. pneumoniae*, 0.1% arabinose for *P. aeruginosa*); growth was monitored by absorbance at 600 nm. Curves are averages of at least two biological replicates.

A compelling feature of CRISPRi is the ease of pooled knockdown library construction, either for defined gene sets or at the genome scale^8^. As a proof of principle, we used Mobile-CRISPRi to construct a 40-member library of selected genes in the opportunistic pathogen, *Enterobacter cloacae* (Supplementary Tables 1 and 2). In the pooled context, each sgRNA functions as a barcode, enabling quantification of each knockdown strain in the pool. To evaluate our pipeline, we performed two different pooled experiments. In the first, all steps from initial cloning to analysis were performed in a pool (Fig. 4a). This revealed that all sgRNA strains were present and had reasonable representation in the pool (31/40 sgRNA counts were within one standard deviation of the median, with a maximum 50-fold difference in representation). In the second, each sgRNA plasmid was constructed individually, and an equimolar mixture of plasmids was used to transform *E. coli* and perform downstream steps (Fig. 4b). This assessed the variability of all steps downstream of cloning and revealed a maximum 2-fold difference in representation. Thus, Mobile-CRISPRi transfer and integration is highly uniform, with essentially all variability derived from the initial cloning step. Further optimization or alternative cloning strategies^8^ may decrease the variability in sgRNA representation.

**Fig. 4.**
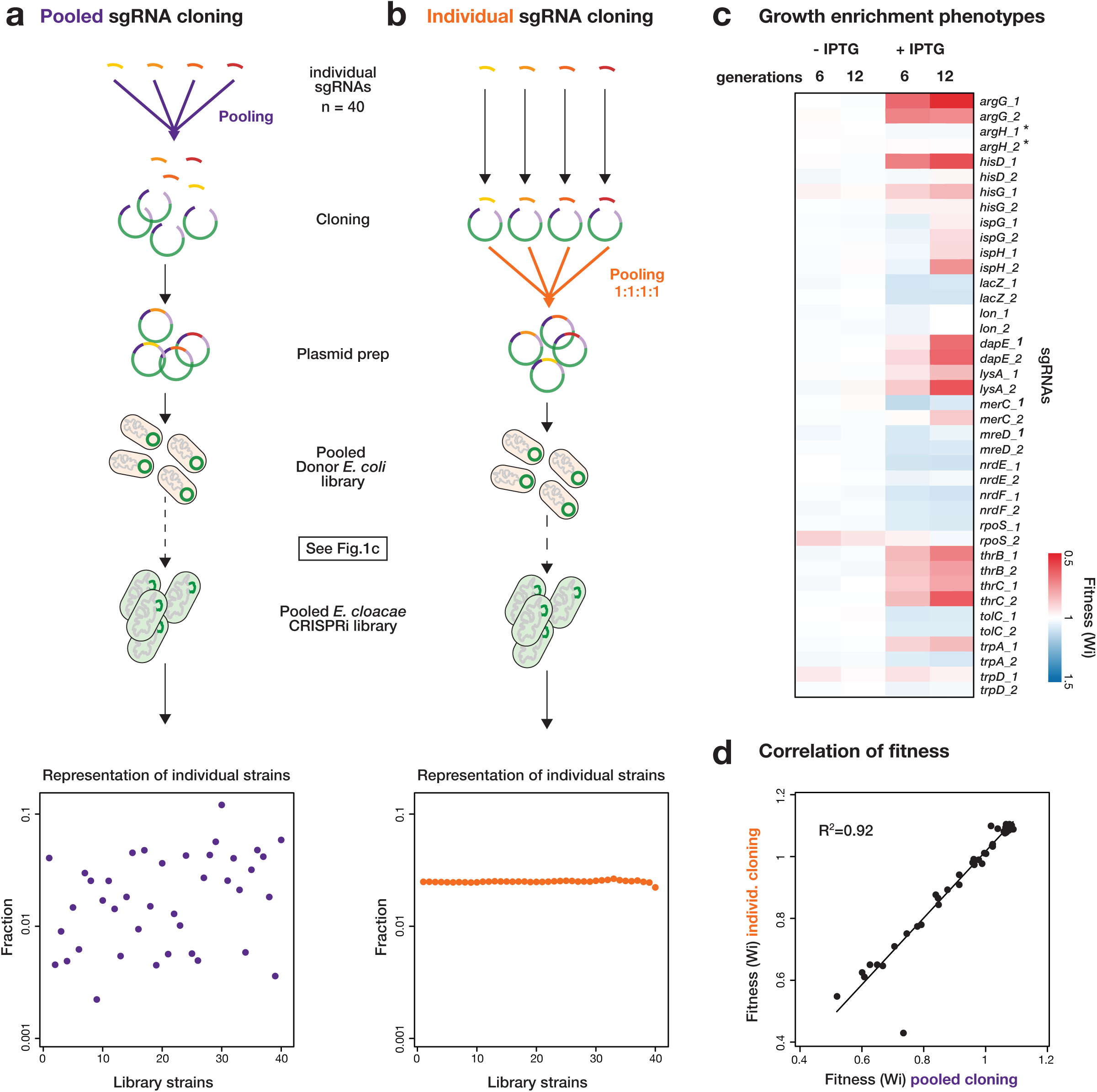
A Mobile-CRISPRi library targeting auxotrophic genes in *E. cloacae*. **a,** Tn*7* Mobile-CRISPRi library construction. sgRNAs were cloned as a pool, transformed, and mated into *E. cloacae*, or **b,** sgRNAs were cloned individually, mixed as a pool with equal representation, and mated into *E. cloacae* as a pool. Representation of individual CRISPRi strains was determined by Illumina sequencing. **c,** Fitness of CRISPRi strains in glucose minimal media after 6 or 12 doublings with or without CRISPRi induction by IPTG, determined from the library constructed by pooled cloning. Asterisks indicate strains had no fitness change in pooled screen but significant decrease in arrayed screen. **d,** Comparison of each strain’s fitness measure by the libraries constructed by pooled cloning (a) or individual cloning (b); a linear fit is shown.

Our library consists of 10 amino acid biosynthesis genes, 4 putative essential genes and 6 well-characterized genes, each targeted by 2 sgRNAs (Supplementary Table 1). For fitness measurements, we grew our library in glucose minimal medium in competition with a 100-fold excess wild-type *E. cloaca*. We measured the relative frequency of each strain in the library after 6 and 12 generations with and without CRISPRi induction to initiate knockdown. Using the fitness calculation of van Opijnen *et al*.^12^, we found that fitness of strains with sgRNAs targeting amino acid biosynthesis and those targeting some putative essential genes decreased, whereas representation of non-essential genes that are unrelated to amino acid biosynthesis remained constant (Fig. 4c; Supplementary Table 2). Fitness for affected strains was more pronounced at 12 doublings than at 6 doublings, suggesting that a larger number of generations was required to dilute out existing protein products. Additionally, both guides generally decreased the fitness of the essential and auxotrophic genes, but with more variability than previously observed^4^, potentially due to cross-feeding of nutrients between strains in the pooled context. Finally, the fitness measurements from the completely pooled construction (Fig 4a), and those in the equal representation library (Fig. 4b, Supplementary Table 2) were highly correlated (R^2^=0.92) indicating that the initial frequency of the strain in the pooled library did not affect the measurement of the fitness (Fig. 4d).

We also screened an arrayed library of individual knockdown strains to confirm the auxotrophy of amino acid biosynthesis gene knockdown strains, finding that their poor growth in minimal medium was suppressed by relevant amino acids (Supplementary Fig. 3 and Supplementary Table 3). Thus, knockdown effects are specific to the targeted gene and do not represent off-target effects of CRISPRi. We conclude that Mobile-CRISPRi enables both pooled and arrayed library construction and straightforward assaying of phenotypes in non-model bacteria.

We anticipate that Mobile-CRISPRi will be a transformative technology for non-model bacteria lacking genetic tools and will facilitate cross-species genetic analysis. The existing Mobile-CRISPRi transfer and knockdown systems are already effective in many organisms, and its modularity makes it straightforward to expand host range (e.g., combining different transfer and integration functions, anti-restriction proteins^22^) and increase knockdown efficacy (e.g., use of “alternative” *dcas9* genes^5,6^). The stability of Mobile-CRISPRi in the absence of selection suggests that it could be a valuable tool for dissecting the genetics of host-microbe interactions in both pathogenic and microbiome contexts, and aid in MOA studies in relevant human pathogens. Interestingly, our approach for transferring CRISPRi mirrors natural transfer of CRISPR systems by transposons related to Tn*7*^23^. We will continue to look to nature for novel approaches to explore the vast landscape of bacterial genetics.

## Figure legends

**Fig. S1 |** The *B. subtilis att*_Tn*7*_ site is not responsible for failure of Tn7 methodology to transfer **a,** Schematic of the transfer experiments. **b,** No transconjugants were obtained from mating a Tn*7* transposon with a *B. subtilis*-compatible kanamycin resistance marker into *B. subtilis*. **c**, Test to determine if the *B. subtilis att*_Tn*7*_ site is responsible for failure of Tn7 methodology to transfer. The *B. subtilis att*_Tn*7*_ site with ~1kb flanking DNA was cloned into a replicative plasmid and transformed into an *E. coli* strain in which integration into the chromosomal *att*_Tn*7*_ site is blocked. E. coli strains with variants of the attTn7 plasmid were used as recipients for a Tn7 transposon containing a kanamycin resistance marker. Kan^R^ transconjugants were obtained from matings with *E. coli* strains containing the *B. subtilis att*_Tn*7*_ site and *B. subtilis* flanking DNA with an *E. coli att*_Tn*7*_ site but were not obtained from an empty vector or *B. subtilis* flanking DNA with a precise deletion of *att*_Tn*7*_, demonstrating that the *B. subtilis att*_Tn*7*_ site is compatible with Tn*7* integration and suggesting that other factors (such as transfer efficiency or transposon gene expression) are limiting.

**Fig. S2 | Mobile-CRISPRi knockdown of native genes in *P. aeruginosa*.** Mobile-CRISPRi was used to target genes involved directly (*phzA1* and *phzM*) or indirectly in pyocyanin biosynthesis (*pqsC*). The loss of blue pigment indicates knockdown of the pyocyanin pathway.

**Fig. S3 | Validation of knockdown-induced auxotrophies by arrayed screen. a,** Construction of an ordered CRISPRi library for *E. cloacae*. sgRNAs were cloned individually, transformed into the *E. coli* donor strain, MFD*pir*. Donor strains were arrayed in 96 well plate and then mating and selection were performed on LB agar plate using a Singer ROTOR robot. **b,** Heat map representation of relative fitness (RF) of 40 *E. cloacae* CRISPRi strains (y-axis) in 7 glucose minimal media conditions with various supplementation (x-axis). Yellow rectangles indicate complementation of auxotrophy of strains by relevant amino acids.

## Methods

### Construction of Mobile-CRISPRi vectors

A complete list of Mobile-CRISPRi vectors can be found in Supplementary Table 4. All plasmids were constructed by restriction enzyme digestion of vector DNA followed by either ligation or NEBuilder HiFi DNA Assembly with insert DNA (all enzymes were purchased from NEB). To generate the Mobile-CRISPRi vectors, the pUC origin of replication in the Tn*7* transposon plasmid pTJ1^24^ was replaced with the R6K γ origin that requires the π protein (encoded by the *pir* gene) for replication, generating pJMP1050, and ensuring that Mobile-CRISPRi vectors cannot replicate in recipient cells. Mobile-CRISPRi “backbone” DNA containing unique restriction sites that flank the cloning modules was synthesized as a gBlock (IDT), and inserted into a pJMP1050 derivative (pJMP1054) that lacked those restriction sites, generating pJMP1055. pJMP1055 served as a base for all Tn*7*-based Mobile-CRISPRi derivatives. New derivatives were constructed by inserting components into the following modules/restriction sites: antibiotic markers/XhoI, reporter genes (e.g., *rfp*)/PmeI, sgRNA promoters and sgRNAs/EcoRI, sgRNA spacers (for creating sgRNA libraries)/BsaI, regulatory genes (e.g., *lacI*)/SmaI, *dcas9* promoters and ribosome binding sites/SpeI, and *dcas9*/SpeI-AscI. To create a Mobile-CRISPRi plasmid that integrates into the ICE *Bs1* element, two ~1kb DNA fragments flanking the *rapI* gene were amplified from *B. subtilis* 168 gDNA and used to replace the Tn*7* transposon ends in a pJMP1055 derivative (pJMP1106), generating pJMP1290. pJMP1290 served as a base for all ICE-based Mobile-CRISPRi derivatives and has the same unique restriction sites listed for the modules above. New sgRNAs were cloned into the BsaI sites of Mobile-CRISPRi plasmids by ligating annealed oligos^8^. Oligos were designed to include overlaps that were complementary to the sticky ends generated by BsaI. Oligos were added to 1X NEB buffer 4 at 5 μM concentration, denatured for 5 min at 95 °C, and then annealed by transferring the reactions to room temperature. Annealed oligos were then diluted 1:20, 2μl of the dilution was ligated to 100ng of BsaI-digested vector for 1hr at room temperature. sgRNAs were designed as previously described^4^.

### Construction of Mobile-CRISPRi strains and mating assays

A complete list of strains used in the study can be found in Supplementary Table 5. Tn*7*-based Mobile-CRISPRi strains were constructed by tri- or quad-parental mating as previously described in Choi *et al*.^15,25^, with several modifications. All Tn*7* matings used MFD*pir*^26^ (a *pir*^+^ strain that is dependent on DAP for growth and contains the RP4 transfer machinery) transformed with either a Tn*7* transposase plasmid (pJMP1039—a derivative of pTNS3^27^ with a spontaneous small deletion upstream of the P_c_ promoter) or transposon plasmid (various pJMP1055 derivatives) as mating donors. Matings with *Acinetobacter baumannii* ATCC19606 required the presence of a third donor strain containing the self-mobilizing RP4 transfer plasmid pRK2013^15^ for unknown reasons. Cultures of the two *E. coli* donor strains (transposon and transposase donors) were grown overnight (~16 hrs) at 37 °C in Lysogeny Broth (LB) + 300 μM DAP (Alfa Aesar B22391) + 100 μg/ml ampicillin. Recipient strains assayed here also grew to saturation in LB after incubation at 37 °C for ~16 hrs. 100 μl of each donor and recipient strain was added to 700 μl of LB and mixed by pipetting. Mixes of donor and recipient strains were pelleted for 2 min at 7000 × *g*, washed twice with 1 ml of LB, resuspended in 30 μl of LB after the final wash, pipetted onto a cellulose filter (MF-Millipore HAWG01300) placed on a pre-warmed LB + 300 μM DAP plate, and incubated at 37 °C for 6 hrs. Filters were then transferred to microcentrifuge tubes containing 200 μl of PBS and vortexed to liberate the cells. Cells were spread onto on media that selects for the Mobile-CRISPRi plasmid and recipient (*e.g.*, LB + kanamycin) without DAP (the absence of DAP will select against donor *E. coli*). Antibiotic concentrations used for selection were: 6 μg/ml (*B. subtilis*) chloramphenicol, 7.5/50 μg/ml kanamycin (*B. subtilis*/*S. aureus*), and 100 μg/ml streptomycin.

ICE-based Mobile-CRISPRi strains were constructed by bi-parental mating as previously described^28,29^ with modifications. New ICE donor strains were generated by transformation of *B. subtilis* with Mobile-CRISPRi integration plasmids using natural competence as previously described^4^. Expression of the ICE anti-repressor, RapI, induces conjugation genes found on the ICE element and promotes excision^28^. ICE excision and the large insert size of Mobile-CRISPRi plasmids resulted in very few transformants. To produce a strain with a stable ICE element in the presence of an IPTG-inducible *rapI* gene that transformed at high efficiency, a *dcas9* gene linked to a chloramphenicol-resistance marker was integrated into ICE—selection for the chloramphenicol marker and the extra homology present in the *dcas9* gene improved transformation efficiency. For mating, one 3 ml LB culture of each donor and recipient strain was grown from single colonies to exponential phase (~2 hrs at 37 °C); donors were grown in LB + 3.25 μg/ml kanamycin to select for ICE retention. Exponential phase cultures were then back diluted to an OD600 of 0.02 and grown until OD600 0.2 before inducing *rapI* expression with 1mM IPTG for 1 hr. 2.5 ml of donor and recipient cells adjusted to an OD600 of 0.9 were mixed with 5 ml of 1X Spizizen salts^30^ and vacuum filtered using an analytical CN filter (Nalgene 145-0020). Filters were transferred to Spizizen agar plates and incubated for 3 hrs at 37 °C. Transconjugants were selected for plating on kanamycin + streptomycin plates as all recipient strains were streptomycin resistant.

### Transfer efficiency assays

Tn7 or ICE mating experiments were carried out in triplicate. Transfer efficiency was calculated by taking the ratio of transconjugants (antibiotic-resistant Dap^+^ colonies for Tn*7* matings, and KanR/StrR colonies for ICE matings) to viable cells (LB colonies for Tn*7* matings, and StrR colonies for ICE matings). For Tn*7* transfer to the *B. subtilis att*_Tn*7*_ site in *E. coli* (Supplementary Fig. 1a), the native *att*_Tn*7*_ site in E. coli K-12 DH10B was occupied by an unmarked Tn*7* to prevent chromosomal transposition, while test *att*_Tn*7*_ sites were cloned onto a chloramphenicol resistant plasmid.

### Mobile-CRISPRi stability assays

Four independently generated isolates of *E. coli* K-12 BW25113 and *B. subtilis* 168 containing Mobile-CRISPRi systems targeting *rfp* were grown to saturation overnight at 37 °C in LB + kanamycin (30 μg/ml for *E. coli* and 7.5 μg/ml for *B. subtilis*) to select for retention of the of the Tn7 or ICE element containing CRISPRi. One ml of each culture was centrifuged at 6000 × *g* for 3 min and washed twice with LB to remove any residual kanamycin. The washed cells were diluted 1:1000 in LB and grown to saturation. The procedure of dilution and growth to saturation was repeated a total of 5 times for ~50 generations of growth. Cells were then serially diluted and plated on selective (LB + kanamycin) and non-selective plates (LB). The ratio between colony counts on LB and LB + kanamycin was used to determine the fraction of cells that retained the Tn*7* or ICE element.

### RFP knockdown assays

RFP knockdown was measured using flow cytometry or a plate reader (for *A. baumannii* and *V. casei*). Flow cytometry was performed by diluting overnight cultures of Mobile-CRISPRi *rfp* knockdown strains 1:10,000 into fresh media (LB for all Proteobacteria and *B. subtilis*, Brain Heart Infusion broth for *S. aureus*) containing CRISPRi inducer (1mM IPTG for all Proteobacteria except *P. aeruginosa*, 1% arabinose for *P. aeruginosa*, and 0.1 μg/ml anhydrotetracycline for Firmicutes) and incubating cultures at 37 °C with rotation until the cultures reached mid-log phase (OD_600_ 0.3-0.6). Cultures were then cross-linked with 1% formaldehyde [final] for 10 min, followed by quenching for 10 min with 0.5 M glycine [final]. Cross-linked cells were then diluted 1:10 in phosphate buffered saline and flowed on a BD LSRII using 610/20 BP filter (PE-Texas-Red fluorochrome). Data for at least 10,000 cells was collected for four independently constructed strain isolates. For *V. casei*, overnight cultures were normalized to 2.0 OD600 and then diluted 1:200 in LB with or without 0.5 mM IPTG. After 6 hours growth post-induction the strains were normalized to 0.2 OD600 and washed once in 1X PBS. The samples were then transferred to a 96well plate (200ul in each well) in triplicate and measured for ds-Red fluorescence (Ex 557nm Em 592nm) using a bottom-read plate reader (Tecan). For *A. baumannii*, overnight cultures were diluted 1:10,000 into fresh LB with or without 0.1 mM IPTG. Cells were grown in a 96 well plate with measurements of OD600 and RFP every 10 min. The values reported reflect the RFP knockdown at mid-log growth.

### Pyocyanin knockdown assays

Strains were grown overnight in Kings Medium A Base (HiMedia M1543) to induce pyocyanin and pyorubin production and 1% arabinose to fully induce *dcas9* expression.

### Antibiotic sensitivity assays

MIC assays were performed using the broth microdilution method as previously described^31^, except that 0.1% arabinose (for *P. aeruginosa*) or 100 μM IPTG (for *E. aerogenes*) was added to induce *dcas9* expression. Growth curves shown in Fig. 3 were set up in exactly the same manner as the MIC assays, except that cultures were grown with agitation in a plate reader (BioTek) for ~16 hrs.

### Construction of Mobile-CRISPRi strains and mating assays

Pooled Tn*7*-based Mobile-CRISPRi libraries for *E. cloacae* were constructed by following the procedure for single gene CRISPRi strain construction with several modifications. Equal concentration of annealed oligonucleotides for each sgRNA (Supplementary Table 1) were pooled and ligated into a BsaI digested plasmid. Ligation product was transformed into an *E. coli pir*+ strain. Colonies on selection plates (LB + 100 μg/ml ampicillin) were collected and resuspended in LB and plasmids were purified from of pooled transformants. Purified pooled plasmids were transformed into donor strain, MFD*pir*. Transformants were collected and resuspended in LB + 300 μM DAP + 100 μg/ml ampicillin + 12.5% glycerol and stored at -80 °C. For comparison, the other donor was prepared by transformation of a pool of individually cloned plasmids with equal concentration. Tri-parental mating and selection were performed as described above and selected colonies of *E. cloacae* CRISPRi strains were collected and resuspended in MOPS salts solution^32^ + 12.5% glycerol and stored at -80 °C after measurement of OD_450_ of stock. In order to prepare inoculum of library to screen fitness of library in minimal media, frozen stock was diluted in glucose minimal medium to OD_450_ of 5 and incubated for recovery for 1 hr. Recovered cell culture was mixed with a 100-fold excess wild-type *E. cloaca*, then diluted to an OD_450_ of 0.01 in 30 ml glucose minimal media with or without IPTG, then grown in 125 ml flasks at 30 °C with shaking (250 rpm). When the culture reached OD_450_ of 0.64, 1 ml of culture was collected for preparation of sequencing library of 6 doubling sample. For 12 doubling sample, this culture was diluted to an OD_450_ of 0.01 in 30 ml and was grown until the culture reached OD_450_ of 0.64. in order to prepare the Illumina sequencing library, genomic DNA was purified using the Qiagen DNeasy Blood & Tissue kit and sequencing region was amplified by PCR using the primers harboring indices for different sampling time and growth conditions. Differentially indexed PCR products were purified by agarose gel electrophoresis prior Illumina sequencing. Sequences of primers used for preparation of sequencing libraries are listed in (Supplementary Table 6). Frequencies of strains in each sample were calculated by dividing the number of reads of sgRNA encoding sequence from each strain by the number of total read and used for calculation of fitness. Fitness was calculated as described in van Opijnen *et al*.^12^ *Wi* = In(Ni(t2)Xd/Ni(ti))/ln((1-Ni(t2))X*d*/(1-Ni(t1))), N(t) is frequency of the mutant in the population at the time points, and d represents the growth of the bacterial population during library selection. We calculated *d* using OD_450_ change. Average fitness from two biological replicates is presented in Supplementary Table 2.

Ordered Tn*7*-based Mobile-CRISPRi libraries for *E. cloacae* were constructed by following the procedure for single gene CRISPRi strain construction with modifications for automation (Fig. S4). Each donor Tn*7*::CRISPRi strains were prepared by transformation of individually cloned plasmids into MFDpir strain and arrayed in 96 well plate. Equal amount of transposase strain was added to each well and pinned to LB + 300 μM DAP + 2% agar plate using a Singer ROTOR robot. Wild-type *E. cloacae* cells arrayed in 96 colony format were pinned to the same plate, which was incubated for 6 hrs. Kanamycin resistant *E. cloacae* CRISPRi strains were selected on LB supplemented with kanamycin two times and stored at -80 °C as a glycerol stock. To screen growth phenotype of each strain, cells were pinned from glycerol stocks onto rectangular LB agar plates in 384-format using a Singer ROTOR robot (four technical replicates on one plate in this screen). For each screen, exponentially growing cells in 384-format were then pinned to defined media plates and incubated for 16 hrs at room temperature to avoid mucoid colony formation. Plates were imaged using a Powershot G10 camera (Canon) when at a time point at which fitness differences were apparent but growth had not saturated. The calculation of RF was carried out as described in Koo *et al*.^32^ with minor modifications. Relative fitness (RF) was measured by the colony opacity of each mutant determined with Iris colony sizing software^33^. The RF of each mutant was calculated as: RF = (average colony opacity of CRISPRi strain)/(average colony opacity of CRISPRi with no sgRNA strain). The average RF calculated from two same media plates is presented in Supplementary Table 3.

## Acknowledgements

We thank Joanna Goldberg and Herbert Schweizer for Tn*7* plasmids, Lei (Stanley) Qi for a plasmid encoding Human codon optimized dCas9, the American Type Culture Collection, KC Huang, Amy Banta, and Paula Welander for strains, Joan Garbarino and Marco Jost for help with flow cytometry, and the Carol Gross and Oren Rosenberg labs for helpful comments. This work was supported by NIH F32 GM108222 (to J.M.P.), US Department of Agriculture National Institute of Food and Agriculture Hatch Project NYC-189438 (to J.E.P.), and NIH R01 GM102790 (to C.A.G.)

## Author contributions

J.M.P., B.M.K., M.M.H., A.D.G, J.E.P., J.N.E., R.J.D., C.A.G., O.S.R. designed the study, J.M.P., B.M.K., R.P., G.E.H., C.C.H., Y.F.I., C.H.S.L. performed experiments, J.M.P., B.M.K., R.P., G.E.H., Y.F.I., J.S.H. analyzed data, and J.M.P., B.M.K., H.O., C.A.G., O.S.R. wrote the manuscript.

